# A high throughput flow cytometry assay for quantifying type 3 secretion system assembly in *Salmonella*

**DOI:** 10.1101/2025.10.25.684470

**Authors:** Jordan Scott Summers, Julie Ming Liang, Nolan Warren Kennedy, Danielle Tullman-Ercek

## Abstract

Recombinant protein production in bacteria is limited by costly cell lysis and multi-step purification. To bypass these limitations, leveraging native bacterial secretion systems to secrete proteins offers a promising alternative. The type 3 secretion system (T3SS) of *Salmonella enterica* serovar Typhimurium can secrete proteins at hundreds of milligrams per liter. While promising, higher titers from this and other natural systems are needed to be relevant commercially.

A major engineering target to enhance protein secretion is the T3SS secretion apparatus. However, efforts to quantify its assembly are hindered by bottlenecks in scalability and complexity. Assembly involves ∼20 structural proteins forming with precise stoichiometry under dynamic regulation. Membrane- and periplasmic-embedded components are difficult to probe without costly, time-intensive methods.

To fully realize the T3SS as a tool for scalable protein production, new tools are needed for rapid and accurate characterization of the assembled apparatus. Here, we establish a high throughput flow cytometry method in *S*. Typhimurium by quantifying the abundance of the needle-bound tip protein SipD as a proxy for T3SS apparatus assembly. We then adapted a super resolution microscopy method, known as Structured Illumination Microscopy (SR-SIM), to visualize the presence of SipD and validate the flow cytometry results. Applying this approach, we revealed how overexpression of key T3SS regulators *hilA* and *hilD* impact assembly, expanded the assay with a secretion-compatible fluorescent reporter to link assembly with secretion, and uncovered how a PrgI variant impacts apparatus architecture. Together, these tools enable rapid insights into T3SS assembly and advance heterologous secretion platform development.

**Importance:** Producing proteins in bacteria often requires expensive and complex steps. This study focuses on *S.* Typhimurium, a bacterium that naturally secretes proteins using a specialized system. We developed a fast and scalable method to measure assembly of the secretion machinery, ultimately accelerating the process of optimizing this system for bacterial protein production. Further, this method provides new mechanistic insights into how the changes to the system impact the ability for the secretion machinery to form. We expect the platform could be adopted to other systems displayed on the bacterial cell surface, including T3SS homologs as well as a wide variety of multi-component membrane transporters.

## Introduction

The secretion apparatus of the Type 3 Secretion System (T3SS) of *S*. Typhimurium is a non-essential pathogenic component that aids in the invasion into host cells. This complex structure composed of over 20 unique proteins spans both the inner and outer membranes, with a distal end needle-like filament and tip complex that plays a pivotal role in sensing host cell contact and initiating the translocation of effector proteins (1). The polymerization of the needle filament monomer, PrgI, and capped with the tip protein, SipD, initiates the host-membrane interaction. This contact induces the secretion of the translocon proteins, SipB and SipC, which insert into the host membrane and form a pore for protein secretion (2). The secreted effector proteins modulate host immunity, cell signaling, and cytoskeletal dynamics, thereby facilitating bacterial uptake, intracellular survival, and replication (3).

In recent years, the T3SS has been repurposed as a scalable platform for heterologous protein production (4), offering a promising alternative to conventional recombinant expression systems. Unlike traditional methods, which often face challenges such as cytotoxicity, protein aggregation, and costly purification processes (5, 6), the T3SS enables direct secretion of target protein into the extracellular environment even in the absence of native T3SS-activating conditions (e.g., it secretes even in the absence of a host cell), simplifying downstream processes and reducing the potential for cytotoxicity (7). Rewiring genetic components, optimizing the extracellular environment, and mutagenizing secretion machinery components for enhanced activity has enabled the production of biomedically relevant proteins into the hundreds of milligrams per liter (8–12). Although conventional intracellular expression followed by cell lysis can yield titers in the gram per liter range, the reduced purification costs associated with secretion-based systems may offset the lower yields. However, to fully harness the potential of bacterial secretion for industrial scale production, further engineering efforts are needed to enhance secretion efficiency and overall system productivity.

Established approaches to increase secretion titer and fine-tune T3SS control include overexpression or knockouts of key T3SS regulators, refactoring to create a minimally viable T3SS, mutations in structural components of the secretion apparatus, or changes to environmental and culturing conditions (8, 10–15). Each strategy is complex, multifactorial, and has potential unintended downstream consequences. Perturbations to the abundance or activity of T3SS regulators can produce dramatic downstream effects, owing to the complex and interconnected regulatory network embedded within the *Salmonella* Pathogenicity Island-1 (SPI- 1). This network integrates an array of intra- and extracellular signals from the environment or culture medium, including pH, bile salts, and osmolarity, which trigger intricate regulatory cascades involving multiple transcriptional regulators (16). Mutations to structural components may alter the physiochemical properties of the needle filament itself, resulting in higher secretion titers, but the interplay between these alterations and T3SS regulation remains underexplored (11, 17). Integration of these synthetic strategies can maximize the potential of the T3SS as a protein production platform, but the combinatorial space of testing these strategies is daunting.

To overcome this challenge, researchers have developed high throughput screening methods to systematically evaluate the effects of genetic and environmental modifications on T3SS secretion titer. Plate-based protein quantification assays have the potential to screen hundreds of regulatory and structural variants in a short time. One effective strategy involves fusing a T3SS-compatible signal sequence to a fluorescent reporter protein, mini Fluorescence Activating Protein (mFAP2a), allowing for real time assessment of T3SS-based protein secretion (18). Additionally, transcriptional activity of key regulators such as *hilD* and *hilA* can be monitored by placing GFP expression under the control of native T3SS promoters and measuring their activation via flow cytometry (8, 19). By combining this transcriptional activity assay with assays for secreted protein titer, such as the mFAP-based assay, upstream regulation can be linked to downstream secretion phenotypes (8, 10–12, 14). These methods have greatly increased the throughput of characterizing activity of T3SS regulators and their impact on protein secretion.

Despite advances in monitoring T3SS promoter activity and secreted protein, the impact of T3SS apparatus abundance or assembly state on secretion titer remains a key outstanding question, and developing high throughput methods to assess the assembly state of the secretion apparatus remains a significant bottleneck to understanding the factors limiting secretion titer. Further complicating this question is the fact that the apparatus is a large, multi-protein complex that assembles across the inner and outer membrane, with many of the components localized in the periplasm or embedded within membrane layers (1). These structural features make it difficult to access and tag many of the individual subunits for antibody or fluorescent labeling techniques. Moreover, the transient and coordinated nature of T3SS assembly complicates efforts to capture its intermediate states. Unlike secretion activity, which can be inferred from extracellular protein levels or reporter fluorescence, the physical assembly of the apparatus lacks a similarly scalable readout.

An attractive and conceptually straightforward approach to assessing T3SS assembly is direct visualization of the secretion apparatus via microscopy. However, the small size of the complex (∼40 nm in base diameter) poses a significant challenge. Conventional light microscopy with a resolution of ∼250 nm cannot faithfully resolve individual apparatuses. To overcome this limitation, several imaging techniques have been employed. Transmission electron microscopy (TEM) and cryo-electron microscopy (cryo-EM) have been instrumental in capturing the secretion apparatus in its membrane-bound state (20), with cryo-EM achieving near-atomic scale resolution (21–23). Cryo-electron tomography (Cryo-ET) has been implemented to directly visualize the secretion needle puncturing host vacuoles (24). Additionally, Zhang and colleagues utilized stochastic optical reconstruction microscopy (STORM) to visualize the spatial organization of two tag-permissive T3SS components, quantifying distances between the membrane embedded protein PrgH and tip protein SipD (25).

While these techniques have provided critical insights into T3SS architecture, they remain inherently low throughput; their time- and cost-intensive requirements, in addition to the extensive technical expertise needed to conduct such experiments, limits their accessibility and scalability. Further, the resource-intensive nature of the technique limits the time points that can be captured of the apparatus, offering limited temporal resolution and making it difficult to observe the progression of apparatus assembly in any detail. Finally, because these techniques operate at the single complex or single cell level, they are impractical for interrogating large libraries of genetic or environmental perturbations in a systematic, high throughput manner. Consequently, insights gained from these methods, while detailed, may not reliably reflect the heterogeneity or overall behavior of the cell population.

To overcome the limitation of low-throughput imaging, we developed a flow cytometry-based method to quantify the T3SS assembly state at the single-cell level with greatly improved throughput and reduced time cost. This approach enables rapid, population-scale measurements while preserving the resolution required to capture cell-to-cell variability. We validated this method by comparing it to direct visualization of the secretion apparatus using SR-SIM and observe strong positive correlations between the measurements gathered via flow cytometry and SR-SIM, confirming the reliability of our approach.

Our method offers several advantages: it is faster, less variable, more cost-effective and scalable, all while maintaining single-cell resolution. Importantly, this method captures the temporal regulatory control of T3SS assembly. We demonstrate this method can report on both the timing and magnitude of assembled apparatuses in response to activation of key T3SS regulators. When combined with a secreted fluorescent reporter protein for expression and secretion, this platform enabled us to deconstruct how different T3SS activation strategies influence secretion rates. Furthermore, we applied this method to a strain harboring a mutation in the structural component PrgI which conferred enhanced secretion titer, and successfully quantified its impact on apparatus assembly.

Together, these results establish a robust, high throughput tool for characterizing T3SS assembly and function. This method not only expands the current toolkit for studying bacterial secretion systems but also provides new mechanistic insights into synthetic activation strategies.

## Methods

### Strains and growth conditions

*S*. Typhimurium strains are derived from ASTE13, a high-secreting derivative of the *S*. Typhimurium strain LT2 (13). Specific strain information can be found in **Table 1.1** in Supplemental Information. All cloning steps were carried out in the *E. coli* strain DH10B. For secretion apparatus assembly, protein expression and secretion experiments, a glycerol stock of the respective strain was streaked out onto a LB-Broth (Lennox) agar plate supplemented with 34 μg/mL chloramphenicol and/or 50 μg/mL kanamycin sulfate where appropriate and placed in a 37 °C incubator overnight to generate single colonies. Single colonies were selected from the plate and used to inoculate a 5 mL LB-L, 0.4% (w/v) D-+- glucose, 34 ug/mL chloramphenicol and/or 50 ug/mL kanamycin (where appropriate) solution in a 24-well block (Axygen). The cultures were then placed in a 225 rpm 37 °C orbital shaker overnight for 12-16 hours. Fresh cultures were prepared with LB-L and subcultured to an optical density at 600 nm (OD600) of ∼0.05. T3SS activation was induced with 100 μM isopropyl β-d-1- thiogalactopyranoside (IPTG) at the time of subculture and grown for respective time courses in a 225 rpm, 37 °C orbital shaker. Cell cultures were then harvested and prepared for subsequent measurements.

### DNA manipulation

All DNA amplification was carried out with either *Q5* or *Phusion* polymerase (New England Biolabs). Primer and plasmid information can be found in **Table 1.2** and **Table 1.3**, respectively. Plasmids verified via Sanger sequencing (Genewiz) were used to transform electrocompetent strains. Successful transformants were identified following selection on LB-antibiotic plates supplemented with the appropriate antibiotic.

### Recombineering

(Adapted from Burdette *et al*., 2021 (10)) Recombineering was performed in *S*. Typhimurium ASTE13 as described by Thomason *et al*. (26). Briefly, a *cat-sacB* cassette conferring chloramphenicol resistance and sucrose sensitivity was amplified from the *E. coli* TUC01 genome using primers with 40 base pairs (bp) of homology 5′ and 3′ to the locus of interest. The desired gene insert was amplified using primers containing the same 40 bp of homology 5′ and 3′ to the locus of interest as used for the *cat-sacB* cassette. PCR was performed with *Phusion* DNA polymerase. *S.* Typhimurium ASTE13 was first transformed with the pSIM6 plasmid encoding the recombination machinery. A first round of recombineering was performed to insert the *cat-sacB* cassette at the locus of interest, and a second round of recombineering replaced the *cat-sacB* cassette with an appropriate DNA product. The replacement DNA was either the wild-type gene with an affinity tag (6x Histidine (His) for *prgI*; 3x FLAG for *sipD*), a combination of a gene variant (i.e., *prgI^S49R^*) with an affinity tag, or a 200 bp double-stranded PCR product containing the first and last 30 bp of the gene to be deleted flanked 5′ and 3′ by 70 bp of homology to the gene locus. Sanger sequencing (Quintara) confirmed the genomic modifications, and the strains were subsequently cured of pSIM6.

### Immunostaining

*Salmonella* cells were harvested and prepared for immunostaining SipD or PrgI on the secretion apparatus first by collecting the equivalent of 1 OD600 of cells in a 2 mL Eppendorf tube. The cell culture was centrifuged at 6000 x *g* for 3 minutes and the supernatant was discarded. The cells were then resuspended in a 4% paraformaldehyde solution in 1x phosphate-buffered saline, pH 7.4 (PBS, Cold Spring Harbor protocol) and gently rocked for 20 minutes at room temperature. After the 20-minute incubation, the cells were centrifuged at 6000 x *g* for 3 minutes and resuspended with 1x PBS, pH 7.4. This wash step was repeated once more then resuspended with 1:1000 dilution mouse-anti-FLAG (Sigma) or a 1:1000 rabbit-anti-6xHis (Sigma) in a 0.5% bovine serum albumin (BSA) 1x PBS solution for 1 hour while gently rocking. The cells were then washed and spun down two more times with 1x PBS, then resuspended with a 1:500 Goat-anti-mouse or 1:500 dilution Goat-anti-rabbit AlexFluor594 fluorescent secondary antibody (Sigma) in 0.5% BSA 1x PBS solution and gently rocked for 1 hour while protected from light. The cells were then washed twice and resuspended with 1 mL of 1x PBS. Cells were either prepped for flow cytometry or, if being prepared for microscopy, were left to incubate with 1 µM of BacLight Green Bacterial Stain (Thermo Fisher) overnight protected from light at room temperature.

### Structured Illumination Microscopy (SIM)

Cells were hard mounted to glass slides by applying cells to a Poly-L-Lysine coated cover slip, adding a drop of ProLong Glass antifade Mountant (Thermo Fisher), inverting the cover slip onto the slide, and leaving to cure overnight while protected from light. Samples were imaged on a Nikon N-SIM system using a Nikon Plan Apo 100x 1.49 NA TIRF objective. Images were captured using Nikon Elements software with a Hamamatsu Flash4 camera. Z-step size was set to 0.100 μm and 15 slices spanning 1.4 µm were taken. 15 images per slice (5 phases, 3 angles) were captured. Images were taken in the 488 channel to visualize the cell bodies (stained with BacLight Green membrane stain) and the 561 channel to detect T3SS apparatus (stained with AlexaFluor 594). Images for each slice were then reconstructed using the Nikon SIM elements software.

### FIJI image analysis pipeline

Sample images were separated into two channels for the detection of the membrane stain and the secondary antibody AlexaFluor 594 conjugate using the FIJI image software (27). Next, the image stack for each channel was compressed into a maximum intensity projection (IP) image. Each channel was then thresholded for cells and puncta, respectively. A selection was created for the masked cell image and overlaid onto a background noise-subtracted puncta image. This overlaid image was used to count the local maxima (prominence >50 to limit false positives) within each region of interest (i.e., cell mask) which yielded a puncta per cell value. Puncta per cell and percent of cells containing puncta values from this image analysis pipeline were analyzed using python matplotlib package (28).

### Flow Cytometry

*Salmonella* samples were diluted to an OD600 of ∼0.03 in 1x PBS and measured using an Attune NxT Flow Cytometer. At least 20,000 events per sample were captured, and the geometric mean of the gated population was used to measure the fluorescence. The arbitrary fluorescence data was converted to a standardized unit (molecules of equivalent fluorescein, MEFL) using BD Sphero Rainbow Calibration Particles (BD Biosciences) according to manufacturer’s guidelines. Data analysis was carried out using FlowJo v10 and visualized using the matplotlib package (28) via Python.

### Western blotting

Whole culture (WC) samples were diluted 1:1 in 4x Laemmli buffer supplemented with 10% BME (final concentration). Supernatant (Sup) samples were harvested by centrifugation of the whole culture for 10 min at 4500 x *g*, then diluting 1:6 in 4x Laemmli buffer with 10% BME (final concentration). Diluted samples were then boiled at 95 °C for 5 minutes, briefly vortexed, then loaded into a 12.5% sodium dodecyl sulfate (SDS) protein gel and separated by performing polyacrylamide gel electrophoresis (SDS-PAGE) at 150 V for 60 minutes. WC and Sup samples were then transferred to an immobilon-P nitrocellulose membrane (Sigma) at 70 V for 30 minutes in a 1x transfer buffer (3 g/L Tris base, 14.4 g/L glycine). The membrane was blocked with a 5% milk 1x Tris-buffered saline with 0.2% Tween-20 (TBS-T) solution, gently rocking for at least 1 hour at room temperature or overnight at 4 °C. The membrane was then incubated with mouse-anti-FLAG (1:6666 dilution) antibody (Sigma) for detection of SipD in a 1% milk TBS-T solution for 1 hour, washed four times with 1x TBS-T (0.20% Tween) solution, then incubated with a goat-anti-mouse-HRP secondary antibody (1:1000) in a 1x TBS-T solution for 30 minutes while gently rocking. The membrane was washed four times with 1x TBS-T, one wash with water, then incubated with the Pierce-ECL western blotting chemiluminescence substrate (Thermo Fisher) for ∼ 30 seconds and imaged on a Workhorse Gel Imager (Azure Biosystems) with 3 minutes of exposure.

### mFAP assay

Fluorescence detection was carried out as previously described (18). Briefly, 100 uL of whole culture samples were combined with 100 uL of 200 uM (5*Z*)-5-[(3,5-Difluoro-4- hydroxyphenyl) methylene]-3,5-dihydro-2,3-dimethyl-4*H*-imidazol-4-one (DFHBI) (Sigma). The secreted fraction was separated from the whole culture by at 4500 x *g* for 10 minutes in a Sorvall Legend XTR centrifuge (Thermo Fisher) using a TX-750 swinging bucket rotor (Thermo Fisher). After centrifugation, 100 µL of the secreted fraction was combined with 100 uL of 100 µM DFHBI. Each DFHBI-sample solution was added to a black flat bottomed 96-well plate and incubated at room temperature while protected from light for 10 minutes. Fluorescence of the samples were measured at 467 nm absorbing and 505 nm emitting using a Synergy H1 plate reader (Agilent) with monochrometer and Gen5 software with a gain of 75. Fluorescence of total expressed or secreted mFAP2a was determined by subtracting the background whole culture or supernatant mFAP2a fluorescence collected from a wild type *Salmonella* strain without a plasmid harboring the *mFAP2a* sequence then subtracting that from experimental values, then OD normalizing. Data was visualized in Python using the matplotlib package (28).

## Results

### The needle apparatus can be quantified and visualized

To determine whether the secretion apparatus could be detected using both flow cytometry and SR-SIM, we fused the C-terminus of SipD with a 3x FLAG epitope tag (hereafter denoted as SipD-FLAG). We used a fluorophore-conjugated secondary antibody for detection and quantification of secretion apparatus assembly at both the single-cell and population levels (Fig.1A). To distinguish cells with and without apparatus, two conditions were tested: (1) overnight cultures containing LB-L supplemented with 0.4% glucose, a known T3SS repressor (29), and (2) cultures grown in an enhanced secretion media that promotes T3SS secretion (10) (LB-ES) without glucose for six hours. Based on the repressive effect of glucose on T3SS, we hypothesized that cells grown in the absence of glucose would assemble T3SS apparatuses, while cells grown in the presence of glucose would not. Following fixation and antibody staining for SipD-FLAG, SR-SIM imaging revealed no detectable SipD-FLAG signal in the glucose-supplemented samples (Fig. 1B). In contrast, cells grown in LB-ES without glucose displayed distinct fluorescent puncta localized near the bacterial membrane (Fig. 1B), consistent with the presence of fully assembled T3SS structures and SipD-FLAG localization at the needle tip.

**Figure 1:**
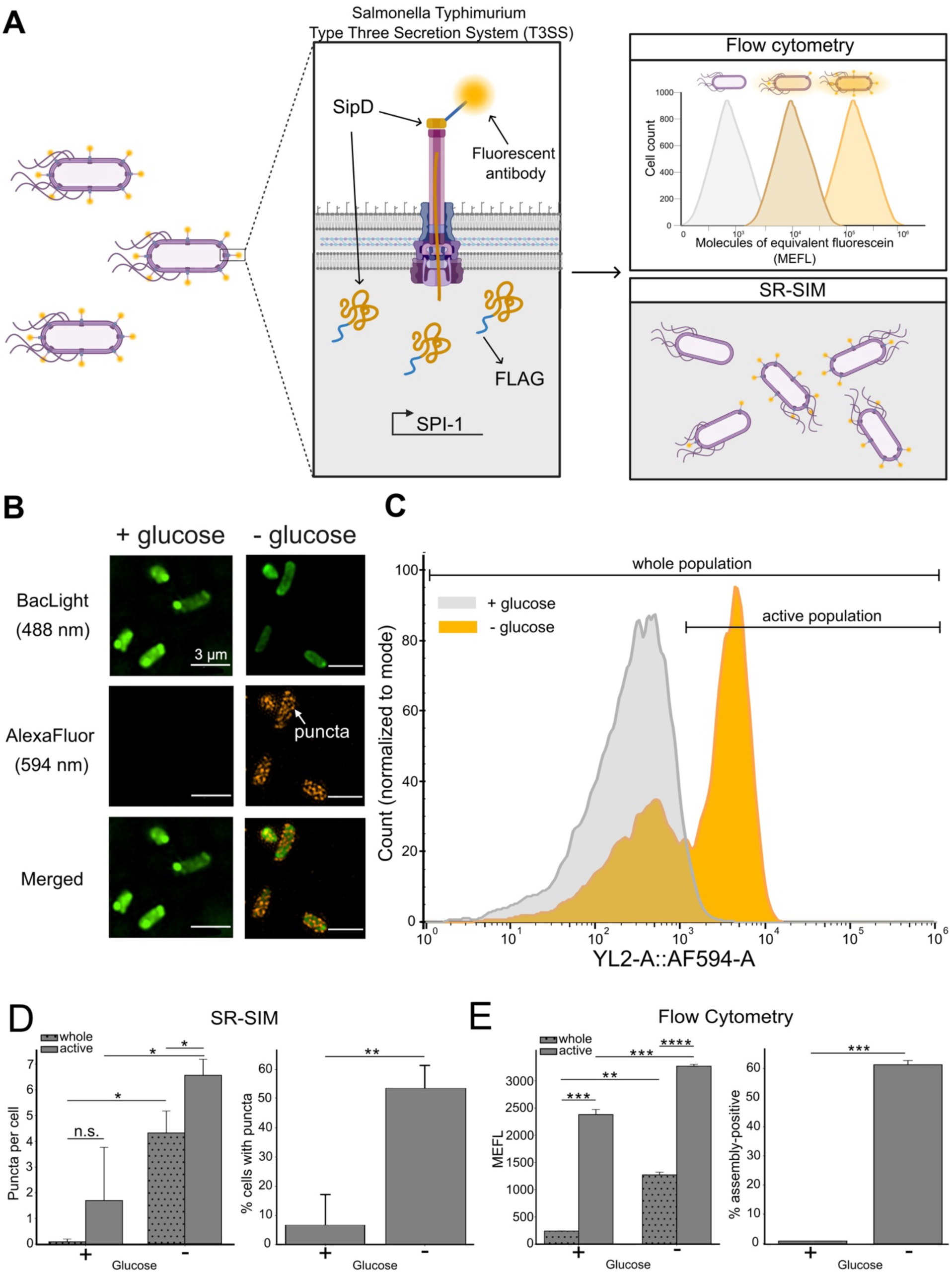
The secretion apparatus can be detected via flow cytometry and SR-SIM. A) The tip protein (SipD) is encoded as a fusion to a 3x FLAG epitope tag. Fixed cells were incubated with a primary α-FLAG antibody, followed by a fluorescently labeled secondary antibody conjugated to AlexaFluor 594. This labeling allows for detection of the SipD-FLAG fusion protein via flow cytometry and SR-SIM, and can be used as a proxy for T3SS assembly. B) SR-SIM images with or without the presence of extracellular glucose, which is known to repress the SPI-1 T3SS. C) Flow cytometry population histograms of the +/- glucose conditions in whole and active (assembly positive) populations. D) Puncta per cell and the percentage of cells with puncta as measured by SR-SIM. E) Representative assembly value (molecules of equivalent fluorescein (MEFL)) and percentage of assembly-positive cells as measured using flow cytometry for cells grown in +/- glucose conditions. Each condition is the average of three biological replicates. Significance was determined using Welch’s unequal variance *t*-test. Significance is denoted by asterisks (*=*p*<0.05, **=*p*<0.01, ***=*p*<0.001, ****=*p*<0.0001). For subfigures B-E, the secondary antibody-fluorophore conjugate is specific for extracellular protein with a FLAG epitope.

The samples were then measured via flow cytometry, revealing distinct fluorescence intensity distributions between samples grown with and without glucose, allowing us to distinguish between the whole population and those containing assembled apparatuses (i.e., “active” population) (Fig. 1C). Consistent with the SR-SIM results, cells cultured in the absence of T3SS inhibitor exhibited higher fluorescence in the active population, indicating increased SipD-FLAG at the cell surface and, by extension, greater T3SS apparatus assembly. These results demonstrate that flow cytometry can be used to reliably detect SipD on the cell surface under activating conditions and further supports the inhibitory effect of extracellular glucose on T3SS assembly.

To ensure that the antibody was specific to the presence of FLAG on SipD-FLAG, we grew cells with and without a genomically-encoded C-terminal FLAG tag fused to *sipD*, and with and without glucose. We then used anti-FLAG western blotting to analyze the proteins from both the whole culture and the secreted spent media fraction of these samples (Supp. Fig. 1A).

As expected, we only detected protein at the expected size for SipD-FLAG when the FLAG tag was present and the system was not repressed. To confirm that our staining method selectively labeled extracellular protein and not intracellular, we expressed the fluorescent reporter protein mFAP2a-FLAG (cytosolic) in a secretion-incompetent Δ*prgI* strain and compared it to our SipD- FLAG strain (Supp. Fig. 1B). We observed no detectable fluorophore signal above background in the strain containing cytosolic mFAP2a-FLAG, confirming the antibody was specific to extracellularly exposed SipD-FLAG.

We then proceeded to extract quantitative and semi-quantitative measurements from SR-SIM and flow cytometry for cells harboring the gene encoding the SipD-FLAG fusion. From SR-SIM, we estimated the proportion of cells containing apparatuses, with ∼52% of cells containing puncta in the absence of the glucose repressor compared to only ∼7% in its presence (Fig 1D, right). Additionally, SR-SIM enabled us to quantify the number of fluorescent puncta per cell as a proxy for the number of assembled apparatuses, observing an average of 6.5 puncta per cell in the active population under glucose-free (active) conditions (Fig. 1D, left). Interestingly, when comparing puncta per cell values in the whole and active populations, (-) glucose cells not only significantly differ from the (+) glucose cells (p<0.05) at the whole population level, their active populations also significantly differ (p<0.05). This indicates T3SS-inducing growth conditions ((-) glucose with LB-ES) drives greater apparatus assembly in populations that display apparatuses. These observations generally agree with activation trends in related species that have the T3SS (e.g., *P. syringae*) (30), insofar that differential expression patterns exist within T3SS “on” subpopulations, representing a potential division of pathogenic labor.

Flow cytometry provided complementary population- and single cell-level data. We calculated the geometric fluorescence intensity for each condition, standardized the mean fluorescence intensity to molecules of equivalent fluorescein (MEFL), and observed a significant difference between each population and subpopulation (p<0.01) (Fig. 1E, left), with the improved statistical significance suggesting flow cytometry may provide more precise measurements than SR-SIM. The ability to collect orders of magnitude more measurements from flow cytometry compared to SR-SIM enables a richer dataset that is difficult to collect purely from micrographs. Flow cytometry also allowed us to estimate the percentage of assembly-positive cells (Fig. 1E, right), yielding similar values to SR-SIM (55.4% and 52.4%, respectively). These results highlight the utility of flow cytometry as a high throughput, single cell resolution method that corroborates microscopy-based measurements.

While SipD-FLAG serves as a useful marker for T3SS assembly, previous studies have shown that secretion titer can increase in Δ*sipD* strains (14, 18), suggesting SipD may not be essential for secretion itself. This makes characterizing combinatorial mutant phenotypes challenging, as single protein tagging strategies are rendered irrelevant in the instance of Δ*sipD* strains. Having established extracellular SipD-FLAG can be reliably detected as a proxy for apparatus assembly, we expanded our approach by targeting an additional extracellular T3SS structural component, PrgI. To this end, we introduced an N-terminal 6x His tag appended to genomic *prgI* and validated its expression and detection using SR-SIM (Supp. Fig. 1C) and flow cytometry (Supp. Fig. 1D).

We compared both protein targets over an eight-hour time course, sampling every two hours. 6xHis-PrgI exhibited a similar temporal trend to SipD-FLAG, though its signal was consistently lower throughout the time course when assessed via microscopy as number of puncta per “active” cell (Supp. Fig. 1C) and flow cytometry as MEFL of the active population (Supp. Fig. 1D). These differences may reflect variations in tag accessibility, antibody affinity, or the relative abundance or exposure of each protein or epitope at the needle filament or tip.

Additionally, we measured the percent of active (e.g., containing puncta in microscopy, or an MEFL above the threshold via flow cytometry) cells over the time course between both methods (Supp. Fig. 1E). Interestingly, we observed that the percentage of active cells after 4 hours are much closer in value for SipD-FLAG and 6xHis-PrgI probing using flow cytometry as compared to SR-SIM at later timepoints (Supp. Fig. 1F). In summary, our results demonstrate that flow cytometry enables high throughput, semi-quantitative assessment of T3SS assembly, with consistent trends observed across different detection platforms and structural targets.

### Assembly of the secretion apparatus is temporally regulated by overexpression of T3SS activators

Activation of the T3SS is a metabolically demanding process with a tightly regulated signaling cascade to ensure proper timing. We have developed several strategies to influence T3SS activation (8, 10–12, 14), with two promising approaches involving plasmid-based IPTG induction of the SPI-1 transcriptional regulators HilD and HilA. HilD sits at the nexus of the SPI-1 regulatory network upstream of *hilA* and directly regulates *hilA* expression, the most downstream and direct activator of SPI-1 (16). Synthetic upregulation of either regulator enhances secretion titers and may also influence assembly state of the secretion apparatus, which would suggest a dual role in both transcriptional and structural control.

To assess how *hilA* and *hilD* influence the assembly of the T3SS apparatus formation, we monitored strains harboring activation plasmids for *hilA*, *hilD*, or no activation plasmid (wild type, WT) over an 8 hour time course using flow cytometry and SR-SIM (Fig. 2A,B). We hypothesized that over expression of these regulators would enhance apparatus assembly, either by increasing the number of assembled apparatuses per cell or by shifting more cells into an assembly-competent state.

**Figure 2:**
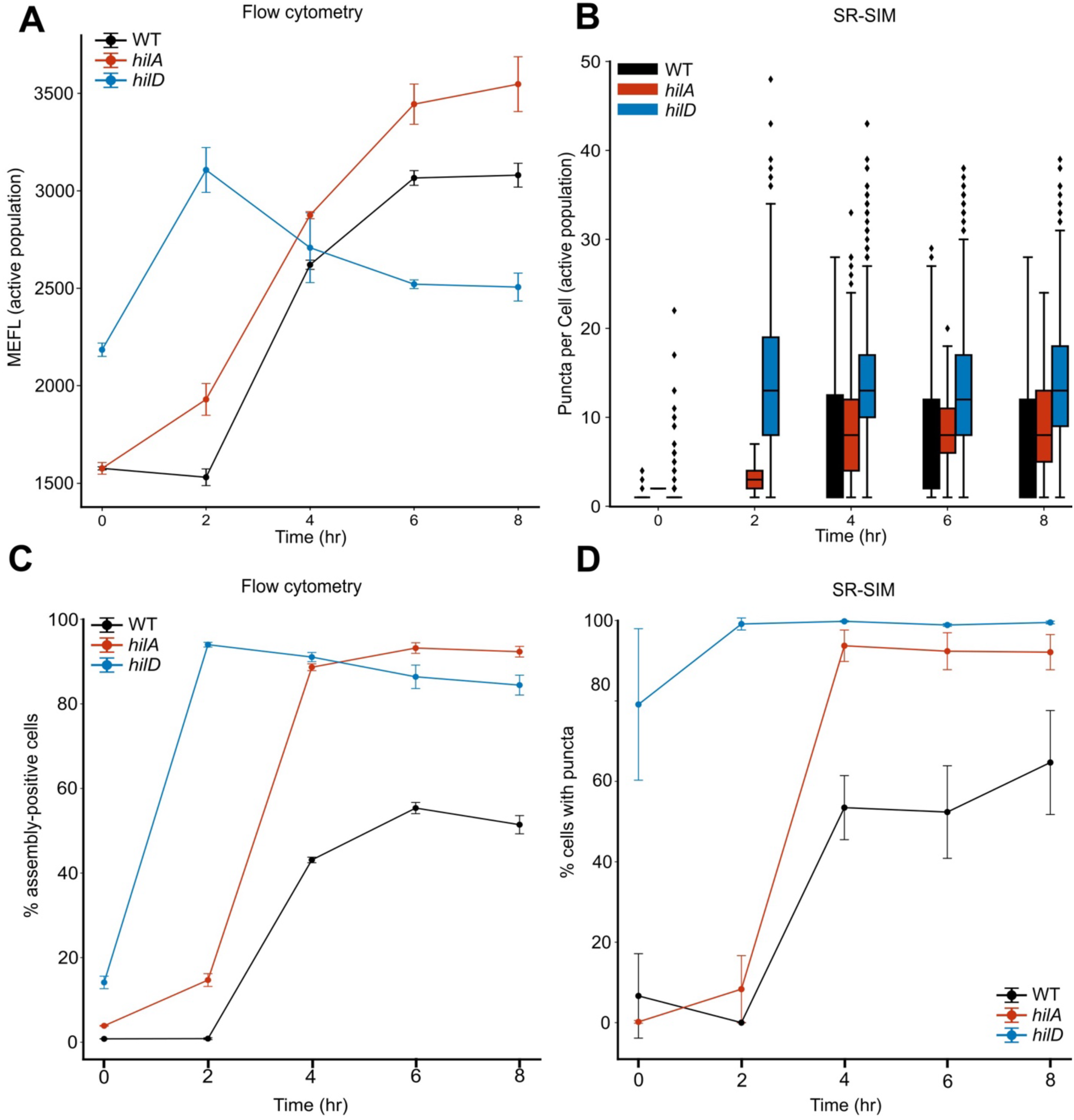
Assembly of the secretion apparatus is regulated temporally and by activation method. A) Assembly of the secretion apparatus measured using flow cytometry (MEFL active population) by probing for the presence of extracellular FLAG-tagged protein in an *S*. Typhimurium strain with a genomic SipD-FLAG tag. B) Assembly of the secretion apparatus measured using SR-SIM puncta counts for the same strains and conditions as assessed in A. C) Percent of assembly-positive cells as measured using flow cytometry from part A. D) Percent of cells containing puncta measured using SR-SIM from part B. For subfigures A-D, a WT *S*. Typhimurium strain, or *S*. Typhimurium strains overexpressing plasmid-encoded *hilA* or *hilD* to activate the T3SS were used for assembly measurements. For subfigures A-D, the secondary antibody-fluorophore conjugate is specific for the primary antibody against extracellular protein with a FLAG epitope.

In the WT strain, we observed a relatively low level of assembly at early time points, followed by a sharp increase between 2-6 hours that plateaued by 8 hours. A similar but amplified trend was seen in the *hilA* activated strain, where assembly increased more rapidly and continued to rise through the 8-hour mark. This sustained increase is consistent with HilA’s role as the most proximal positive regulator of T3SS structural genes, suggesting that its overexpression enhances apparatus formation by directly upregulating transcription of the *prg* and *inv* operons that form the structural components of the apparatus (12, 31). In contrast, *hilD* activation resulted in high assembly levels even at the 0-hour time point, indicating the greater abundance of HilD may bypass early repression mechanisms, such as the presence of glucose in the 0-hour culture, and initiate apparatus formation more rapidly or rescue the apparatus-positive phenotype. This early activation is likely due to HilD’s capacity for auto-activation (32, 33), and, similarly to HilA’s impact on structural promoter activity, may explain the sharp increase in assembly observed at 2 hours. However, unlike the *hilA* over expression and WT conditions, the assembly trend in *hilD*-activated cells gradually decreased over time. This tapering may reflect a post-transcriptional regulatory feedback mechanism mediated by the 3’ untranslated region (3’UTR) of *hilD* (34). The exogenously induced HilD may be autoactivating its own genomic expression, which increases the amount of genomically transcribed *hilD* containing the 3’UTR. This may promote *hilD* mRNA turnover (34), resulting in less abundant HilD and decreased T3SS activity as the system stabilizes over the time course.

Both activation strategies significantly increased the proportion of cells exhibiting puncta, as detected by both flow cytometry and SR-SIM (Fig. 2C, D), with a strong positive correlation between the two methods (Supp. Fig. 2A). These findings support the idea that transcriptional activation of T3SS regulators not only elevates gene expression but also drives physical assembly of the secretion apparatus. Our data provides direct evidence that this transcriptional upregulation translates into enhanced levels of apparatus assembly in the population.

Previous research has shown synthetic activation of the T3SS can increase the proportion of cells within a population to become T3SS active (8, 12). Building on this, we hypothesized that activation strategies over expressing *hilA* or *hilD* not only elevate population assembly levels but also modulate single-cell characteristics of T3SS assembly, potentially leading to altered numbers of apparatuses per cell within the T3SS-positive population. To test this, we compared strains overexpressing *hilA* or *hilD* to WT, analyzing both the whole and active populations forming apparatuses averaging across the entire time course sampled. Our results indicate that both activation methods increase the puncta per cell (Supp. Fig. 2B, top) and the MEFL (Supp. Fig. 2B, bottom) across the whole population compared to WT. When focusing on the active population, *hilD* overexpression confers a significantly higher puncta per cell and MEFL value compared to WT (p<0.01). The exception is the SR-SIM data in the *hilA* active population, which did not show statistical significance, likely due to higher variability in image-based quantification. Nonetheless, both SR-SIM and flow cytometry show strong agreement overall and indicate potential future activation strategies to push not only more cells into an assembly competent state, but to maximize the subpopulations containing higher assembly values (i.e., apparatuses per cell).

Our findings suggest that synthetic T3SS activation may play dual roles for protein secretion by increasing the proportion of cells with assembled apparatus and by boosting the number of apparatuses per cell. This shift appears to bypass cell density-dependent T3SS regulation (35, 36), as *hilA* and *hilD* activation results in higher MEFL and puncta counts even at lower OD600 compared to WT (Supp. Fig. 2C). This decoupling of population control mechanisms and T3SS activity highlights the potential of synthetic strategies to optimize T3SS output by tuning population-level and single-cell assembly dynamics, and the ability of this assay to assess these dynamics. Insights gained through this flow cytometry-based method are key in optimizing recombinant protein production via the T3SS.

### Characterizing the relationship between assembly and protein secretion

Activation of the T3SS is tightly regulated by environmental cues such as host cell contact, salt and osmolarity, and interbacterial signals, which initiate apparatus assembly and effector secretion. In repurposing T3SS for recombinant protein export, simply fusing secretion signals or chaperone binding domains to a target protein often results in incomplete secretion of intracellular protein, potentially due to partial apparatus assembly or native regulatory suppression. Our flow cytometry method to measure apparatus assembly lets us combine this tool with other methods that can measure protein expression and secretion, delivering a holistic assessment of these various aspects of T3SS activation. To that end, we measured assembly via SipD-FLAG in conjunction with a compatible secreted fluorescent reporter protein, mFAP2a, enabling dynamic monitoring of apparatus formation alongside protein expression and secretion (18). This dual assay framework allows us to identify bottlenecks in secretion and test whether synthetic activation strategies can bypass native regulatory constraints.

We measured assembly via the flow cytometry method described above, and expression and secretion of our fluorescent reporter, mFAP2a, via a microplate reader, for our WT strain as well as using strains overexpressing *hilA* and *hilD* over 8 hours (Fig. 3A). Generally, we observe that over time each condition produces increasing assembly, expression, and secretion values. Assembly follows similar characteristics to those observed in Figure 2, and the highest amounts of secreted protein are detected at the end of the time course. As expected, higher assembly (MEFL) values precede the detection of expressed and secreted mFAP2a, indicating a controlled temporal mechanism of T3SS activity: assembly of the secretion apparatus occurs first, followed by expression and secretion of select effectors/chaperones soon after.

**Figure 3:**
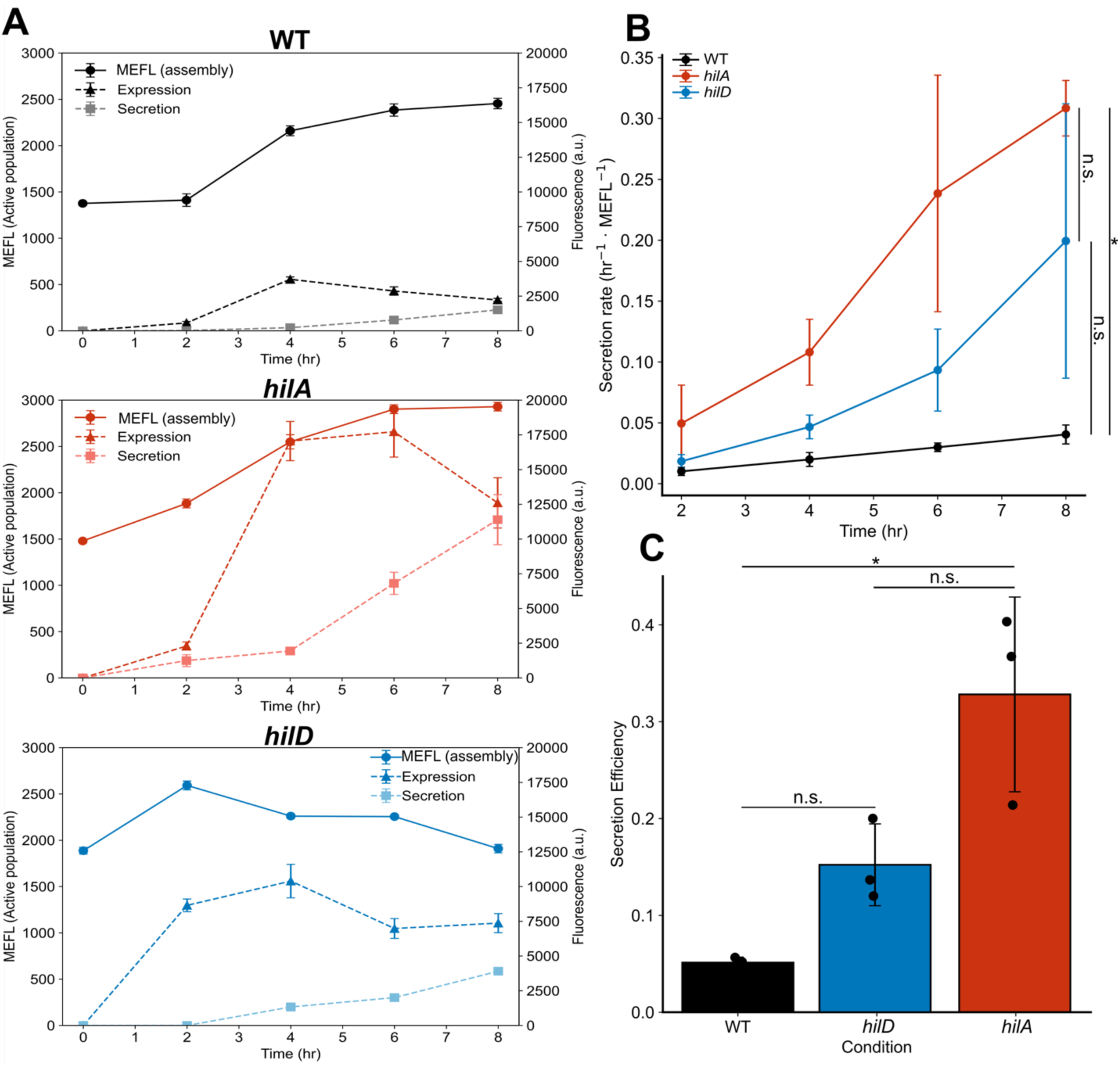
Characterization of assembly, protein expression and secretion. A) Temporal relationship between assembly of the secretion apparatus, and expression and secretion of a target protein, mFAP2a. The left y-axis defines assembly of the active population (MEFL), using flow cytometry to measure the presence of extracellular FLAG-tagged protein in an *S*. Typhimurium strain with a genomic SipD-FLAG tag. The right y-axis defines fluorescence in arbitrary units (a.u.) of the expressed and secreted protein, mFAP2a, which is under the control of a plasmid-based native T3SS *PsicA* promoter. Fluorescence of expressed or secreted mFAP2a was calculated by subtracting background fluorescence from a WT *S*. Typhimurium strain (lacking the mFAP2a expression plasmid), normalizing to optical density, and measured using a microplate reader. B) Rate of secretion under different T3SS activation conditions. Secretion rate was determined by normalizing to the highest experimental value for mFAP2a fluorescence and assembly (MEFL), respectively. Significance was determined by performing a one-way ANOVA. Significance is denoted by asterisks (*=*p*<0.05, **=*p*<0.01, ***=*p*<0.001, ****=*p*<0.0001). C) Secretion efficiency rate per unit assembly averaged over an 8 hour time course. For all subfigures, each condition is the average of three biological replicates. Significance was determined using Welch’s unequal variance *t*-test. Significance is denoted by asterisks (*=*p*<0.05, **=*p*<0.01, ***=*p*<0.001, ****=*p*<0.0001). For all subfigures, a WT *S*. Typhimurium strain, or *S*. Typhimurium strains overexpressing plasmid-encoded *hilA* or *hilD* to activate the T3SS were used for assembly measurements. The secondary antibody-fluorophore conjugate is specific for the primary antibody against extracellular protein with a FLAG epitope.

The combination of the two high-throughput methods is a powerful tool for studying assembly and secretion dynamics, particularly those that are important for downstream applications of T3SS mediated protein production, such as secretion efficiency and secretion rate. While these metrics have been quantified in the context of total protein made and secreted protein, these methods lacked the ability to quantify efficiency and rate with respect to secretion apparatus-mediated protein secretion (i.e. protein secreted per unit assembly). From these methods, we were able to calculate the rates of secretion through the apparatus and found that *hilA* overexpression confers significantly higher secretion rates compared to WT (p<0.05) (Fig. 3B). Additionally, the *hilA* overexpression strain more efficiently secretes protein than WT over the entire time course (Fig. 3C), indicating that synthetic activation increases secretion titer using a combination of assembly and expressed protein dynamics. We also note that assembly and secretion exhibit positive correlation (Supp. Fig. 3), further emphasizing the close relationship between assembly and secretion dynamics. It is possible the observed secretion rate and efficiency gains seen with synthetic T3SS activation are the result of both higher assembly and expressed protein; while more protein is being expressed, the available conduits for protein trafficking contribute to rapid and efficient protein export.

### Flow cytometry reveals mechanisms of structural mutants on protein secretion and assembly

We have shown our flow cytometry method can faithfully measure assembly across various conditions, particularly those that directly activate the T3SS, and these measurements corroborate well with our SR-SIM measurements. Further improvements to increasing secretion titer may lie beyond upregulation of key activators or deletion of inhibitors. As the main physical barrier in protein translocation, improvements made to the structural components that make up the secretion apparatus via mutation serve as an attractive area to optimize protein secretion.

Numerous studies have explored the impact of mutations of structural components on the ability of Gram-negative bacteria to secrete protein into host cells or successfully form an apparatus (23, 37, 38). In-depth mutagenesis libraries show the needle filament protein PrgI can be mutated at select residues to confer higher secretion titers (11). Of these high-secreting variants, a serine to arginine mutation at position 49 (S49R) mutation may be involved in inter-subunit interactions, leading to more favorable multimerization of PrgI as it forms the secretion needle or by translocating target proteins in the unfolded state more efficiently. However, these hypotheses have not been explored fully due to a lack of reliable methods to concurrently assess assembly and secretion.

We attempted to answer this question by measuring the assembly state of the mutated PrgI-S49R apparatus using our flow cytometry approach, again probing for the presence of SipD-FLAG on the cell surface. We were initially surprised to see that throughout the time course, the assembly value for the PrgI-S49R mutant remained no higher than background while the WT’s assembly value progressed in a manner consistent with our previous experiments (Fig. 4A). Further complicating our observation was the increased amount of secreted reporter protein, mFAP2a, in the PrgI-S49R strain compared to WT (Fig. 4B), as expected based on prior reports (11).

**Figure 4:**
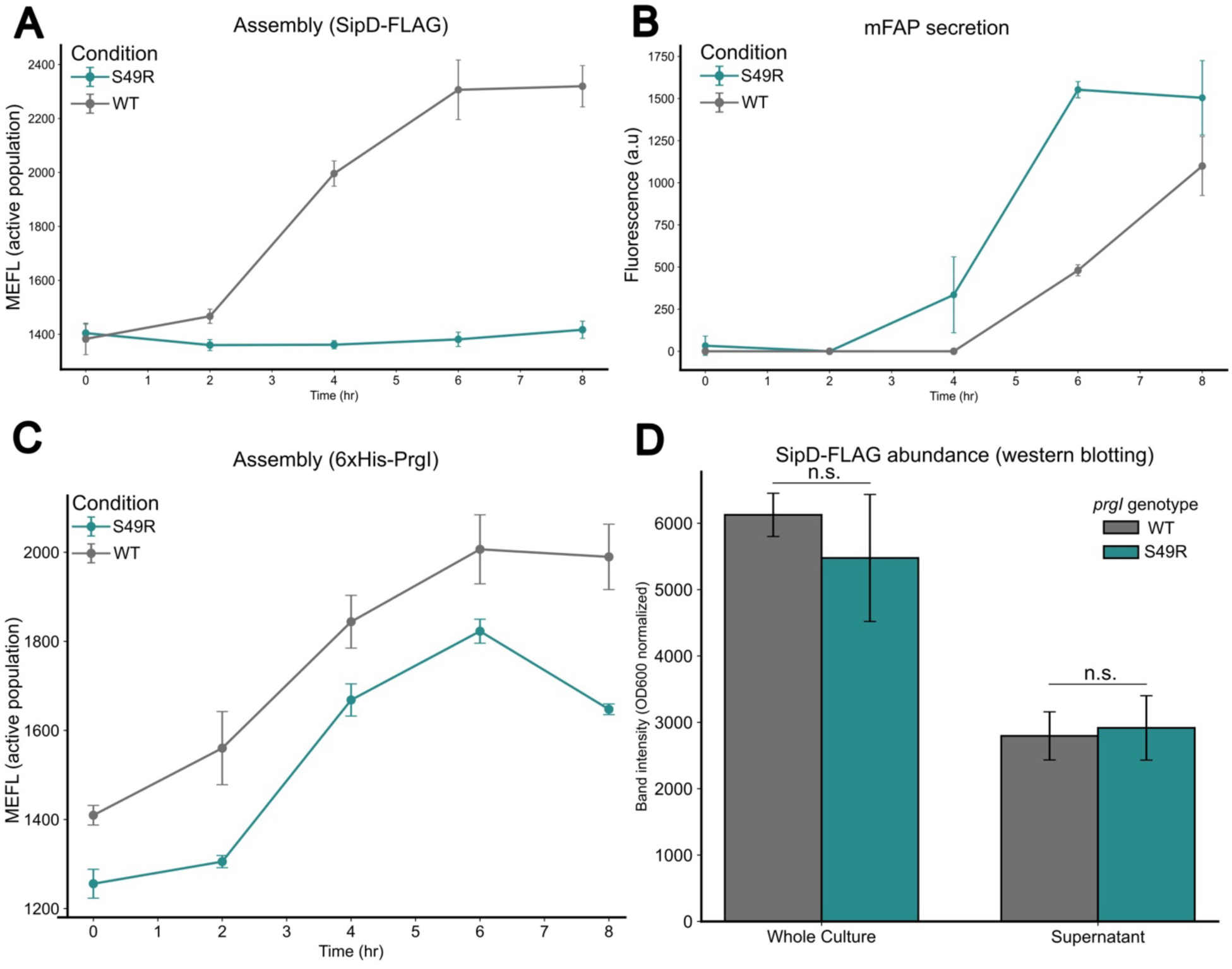
Assessing the impact of the S49R mutation in PrgI (6xHis-PrgI-S49R) on assembly of the secretion apparatus and protein secretion. A) Assembly of the secretion apparatus as assessed by flow cytometry (MEFL active population) over 8 hours by probing for the presence of extracellular FLAG-tagged protein in WT PrgI and S49R mutant PrgI strains. B) Secretion of mFAP2a over 8 hours in both PrgI WT and S49R strains. Fluorescence of the secreted mFAP2a was calculated by subtracting background fluorescence from a WT *S*. Typhimurium strain (lacking the mFAP2a expression plasmid), normalizing to optical density, and measured using a microplate reader. C) Assembly (MEFL, active population) over 8 hours in PrgI WT and S49R mutant strains by probing for extracellular 6xHis-PrgI using flow cytometry. A fluorescent secondary antibody with a conjugated AlexaFluor 594 fluorophore is bound to the primary α- 6xHis antibody on 6xHis epitope fused to PrgI. D) Semi-quantitative band intensity reporting on the abundance of SipD-FLAG in the whole culture and supernatant in either WT or S49R PrgI strains cultures sampled from the 8 hour timepoint. Band intensities were quantified using SDS- PAGE protein separation, probing of SipD-FLAG via western blotting, and quantified intensity using Fiji (27). For all subfigures, each condition is the average of three biological replicates. Significance was determined using Welch’s unequal variance *t*-test. Significance is denoted by asterisks (*=*p*<0.05, **=*p*<0.01, ***=*p*<0.001, ****=*p*<0.0001). For subfigures A and C, the secondary antibody-fluorophore conjugate is specific for the primary antibody against extracellular protein with a FLAG or 6x His epitope, where appropriate.

These observations led us to question if SipD-FLAG was present in the secreted fraction or if the S49R mutant was disrupting SipD-PrgI interaction, as previous studies have reported increased secretion titer in the presence of non-PrgI-binding SipD mutants (14). To explore this hypothesis, we probed for the presence of extracellular PrgI in both WT and PrgI-S49R strains with the N-terminal 6xHis tag and find that while assembly is slightly lower than WT throughout the time course, the trends are similar (Fig. 4C). These results indicate the 6xHis-PrgI-S49R assembly remains intact while the SipD interaction with PrgI is disrupted. The reduced MEFL value for the 6xHis-PrgI-S49R strain suggests that the epitope tag is less accessible for detection by flow cytometry.

Finally, we wanted to determine the abundance of SipD-FLAG in the extracellular space in the presence of the 6xHis-PrgI-S49R, as the point mutation may impact the ability to secrete, which also would lead to a low signal when measuring a SipD-FLAG-bound fluorophore on the apparatus. We collected samples of the whole culture and supernatant and probed for the presence of SipD-FLAG in both WT and 6xHis-PrgI-S49R conditions using western blotting and observed no significant change in the abundance of expressed or secreted SipD-FLAG protein (Fig. 4D).

These findings suggest SipD-FLAG is expressed and secreted in our 6xHis-PrgI-S49R mutant strain in a manner similar to the strain harboring the non-mutated PrgI, but the S49R mutation disrupts the interaction between the two subunits of the secretion apparatus. These observations highlight the utility of our flow cytometry assay, permitting detection of more than one component of the secretion apparatus to provide relevant information on its assembly status. Our method also illuminated the mechanism by which promising candidate mutant strains bypass native physical constraints of assembly to produce higher secretion titer and point to future optimization targets for T3SS mediated protein production.

## Discussion

To more rapidly characterize and identify optimization bottlenecks in repurposing native bacterial machinery, reliable, high-throughput methods to measure multiple components of the T3SS and other secretion systems are crucial to develop. The incorporation of scalable methods to quantify secretion systems serves the greater goal of using secretion systems for myriad purposes. Our implementation of this flow cytometry method substantially reduced the time and cost associated with data collection while simultaneously decreasing the variability compared to SR-SIM (Supplemental Table 1).

Our method further characterizes the mechanisms that drive protein secretion from the perspective of secretion apparatus assembly. Not only can these insights provide a greater understanding of assembly, but the disassembly of the apparatus as well, highlighting important factors that may result in lower secretion titer, like in the case of *hilD* overexpression. This method, in conjunction with other high throughput methods, such as GFP-promoter fusions and T3SS compatible fluorescent reporters, can provide a comprehensive understanding of the major contributors of T3SS activation, how this relates to protein secretion, and further expands our understanding as to why overexpression of key T3SS regulators result in higher secretion titer, thus deepening our ability to understand and reengineer the system.

Notably, our flow cytometry method provided greater resolution in identifying how overexpression of key T3SS activators can bypass regulatory mechanisms that typically suppress T3SS activity. Overexpression of either *hilD* or *hilA* led to significantly elevated assembly values compared to WT at similar time points (Fig. 2A). Interestingly, at earlier time points, these overexpression strains exhibited lower cell densities than WT but still showed markedly higher assembly levels (Fig. 2A, Supp. Fig. 2C). Further, *hilA* overexpression displayed increased assembly rates even into the 8-hour time point, suggesting assembly plateaus later than the sampled time range. This suggests T3SS activation via *hilA* or *hilD* overexpression may be less dependent on population-level regulatory signals (39). These findings point to the need for further exploration of the relationship between cell density-mediated cellular communication mechanisms, such as quorum sensing (40), and physiologically divergent T3SS activation strategies like plasmid-based activator overexpression.

Our method greatly expands experimental sample sizes compared with current “gold standard” methods like super-resolution microscopy to further isolate and more thoroughly measure distinct populations, such as T3SS “off” and “on” populations. Not only are these populations distinct from one another, the “on” populations significantly change in *hilD* and *hilA* overexpression strains. We observe a significant difference in MEFL and puncta per cell when populations only containing fully assembled apparatus are present (Supp. Fig. 2B). This suggests that overexpression of these key activators not only changes the percent of the population that have apparatuses but also changes the number of apparatuses per cell, which agrees with similar SPI-1 subpopulation transcriptional and growth trends (41). This introduces an important balance for future optimization efforts to explore. Controlling the optimal amount of apparatus per cell, percentage of the population that contains apparatus, and the timing of POI expression will be a multifactorial challenge. For example, diverting a majority of cellular resources to T3SS activation and apparatus assembly (41) may result in poor protein expression and less secreted protein. Our flow cytometry-based method for assessing assembly provides a deeper biological context and can aid in developing optimal strains.

While we have demonstrated this method is a powerful addition to the toolkit for T3SS reengineering purposes, we envision some iterations for further scaling and expanding this method’s capabilities. Using multiplexed labeling can facilitate the reporting of several structural components concurrently, and an ideal reporter system would implement a secretion compatible fusion tag that is inherently fluorescent and eliminates the need for antibody incubation steps. Incorporation of a tag that fluoresces upon binding to the needle tip could also provide a real time reporter straight from the cell culture, thus significantly reducing the downstream processing required to measure assembly via flow cytometry. Ultimately, establishing and refining this method will be essential for developing a robust, reliable, and scalable platform for T3SS-mediated recombinant protein secretion.

## Acknowledgements

We would like to acknowledge Dr. Constadina Arvanitis and Dr. David Kirchenbuechler at the Center for Advanced Microscopy at Northwestern University for their microscopy expertise. We thank Dr. Kevin Metcalf, Dr. Han Teng Wong, and Daniel Payan for contributing the *Salmonella* ASTE13 strains.

JSS and DTE were supported by the National Science Foundation (award numbers BBE- 1706125; DMR-2324252). JSS was also supported by the National Institutes of Health Training Grant (T32GM008449) through Northwestern University’s Biotechnology Training Program.

## Author Contributions (CRediT)

**Jordan Scott Summers:** Conceptualization; methodology; investigation; writing – original draft; validation visualization; project administration; formal analyses; data curation; funding acquisition; writing – review and editing; resources. **Julie Ming Liang:** Conceptualization; methodology; project administration; writing – review and editing; resources. **Nolan Warren Kennedy:** Conceptualization; project administration; writing – review and editing. **Danielle Tullman-Ercek:** Conceptualization; funding acquisition; investigation; methodology; validation; writing – review and editing; software; formal analysis; project administration; data curation; supervision; resources.

## Notes

### Competing Interest Statement

Danielle Tullman-Ercek and Julie Ming Liang have a financial interest in Opera Bioscience, which is commercializing bacterial protein production and secretion. The conflict of interest is reviewed and managed by Northwestern University in accordance with their conflict of interest policies.

