## Supplemental Figures for "A high throughput flow cytometry assay for quantifying type 3 secretion system assembly in *Salmonella*"

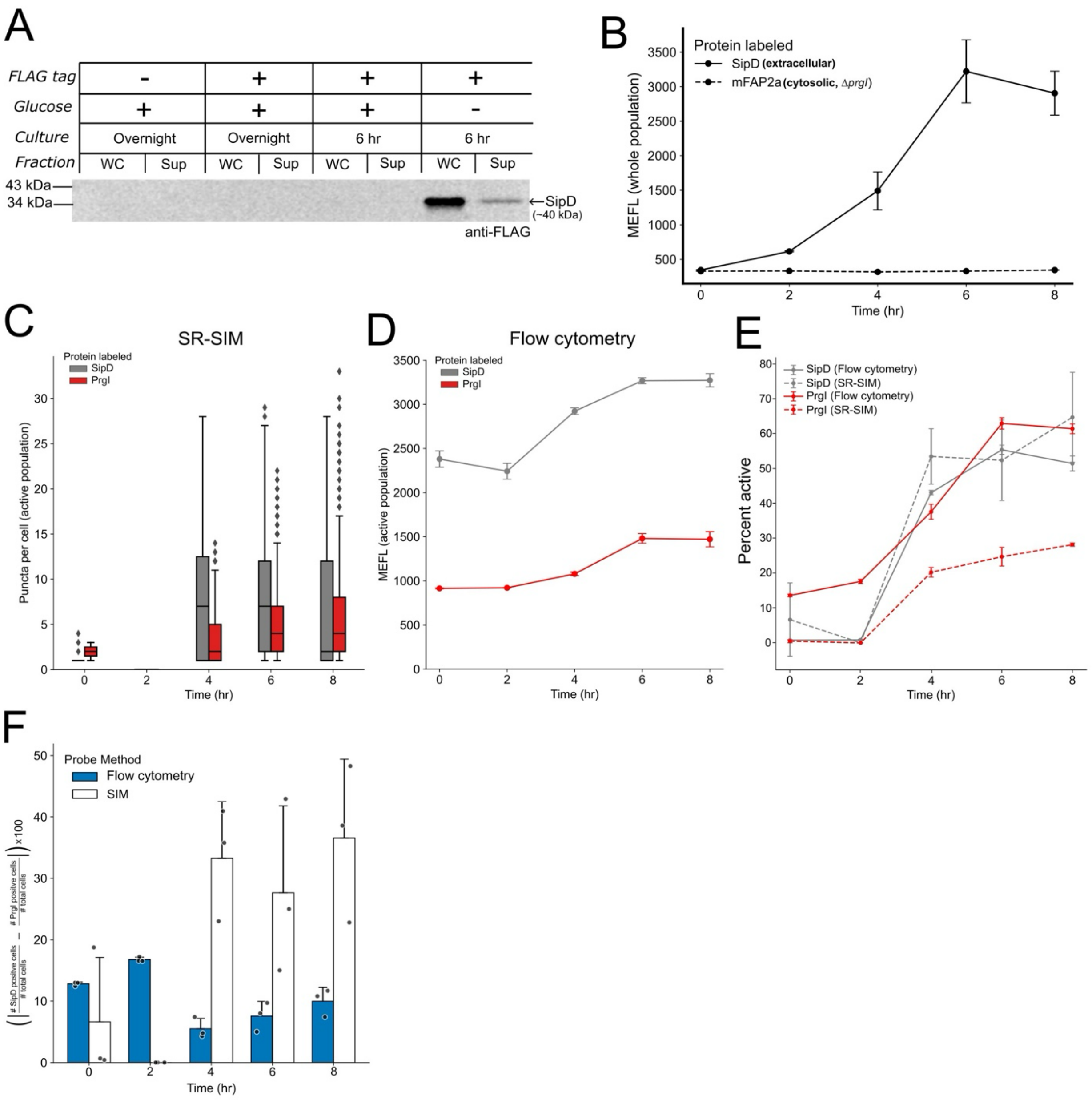


**Supplemental Figure 1:** Establishing detection of SipD-FLAG and 6xHis-PrgI. A) Representative western blot of whole culture (WC) and supernatant (Sup) fractions probing for FLAG epitope fused to the C-terminus of SipD (expected size ~40 kDa) under specific culturing conditions. B) Two *S*. Typhimurium strains were measured via flow cytometry (i.e., assembly (MEFL)) for the presence of extracellular FLAG-tagged protein: (1) A genomic SipD-FLAG tagged strain, and (2) a secretion-incompetent Δ*prgI* strain harboring a plasmid containing mFAP2a-FLAG. C) Puncta per cell (active population) counts for SipD-FLAG and 6xHis-PrgI strains over an 8 hr time course assessed using SR-SIM. D) Assembly (MEFL, active population) assessed over an 8 hr time course via detection of either SipD-FLAG or 6xHis-PrgI using flow cytometry. E) Percent active (assembly positive) cells assessed over an 8 hr time course via detection of either SipD-FLAG or 6xHis-PrgI using flow cytometry or SR-SIM. F) Absolute % difference between SipD-FLAG and 6xHis-PrgI positive cells from (E), measured by flow cytometry or SR-SIM. For subfigures B-F, the secondary antibody-fluorophore conjugate is specific for extracellular protein with a FLAG or 6xHis epitope, where appropriate.

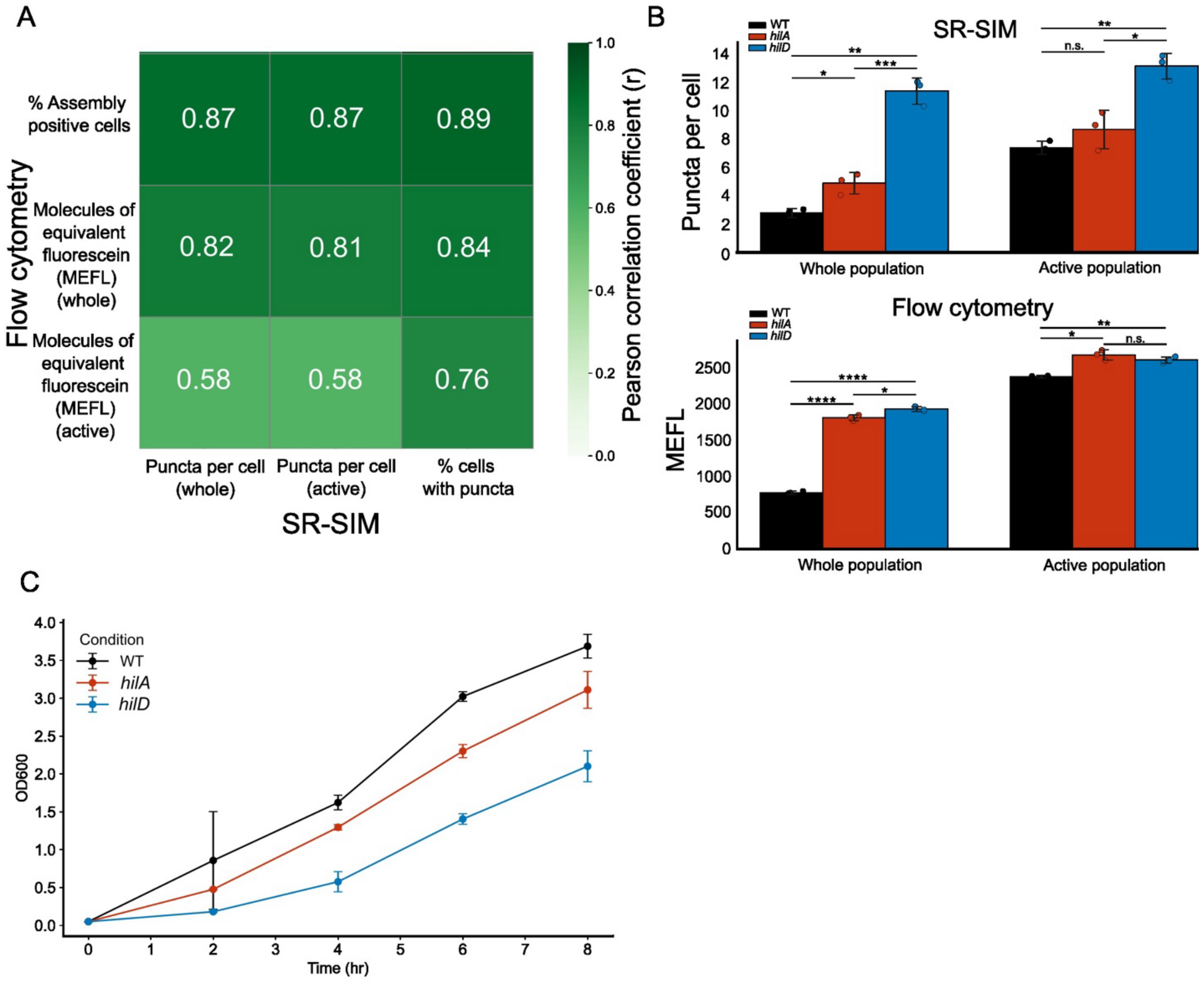


**Supplemental Figure 2:** Comparisons of flow cytometry and SR-SIM measurements and growth characteristics. A) Pearson correlation coefficient (r) heatmap between the flow cytometry and SR-SIM derived measurements. B) Average puncta per cell (SR-SIM) and MEFL (flow cytometry) measurements for a WT *S*. Typhimurium strain, or *S*. Typhimurium strains overexpressing plasmid-encoded *hilA* or *hilD* to activate the T3SS, assessed over an 8 hr time course. Significance was determined using Welch's unequal variance t-test. Significance is denoted by asterisks (*=p<0.05, **=p<0.01, ***=p<0.001, ****=p<0.0001). C) Optical density assessed via measuring absorbance at 600 nm ($OD_{600}$) over an 8 hr time course for WT, *hilA* and *hilD* activation conditions.

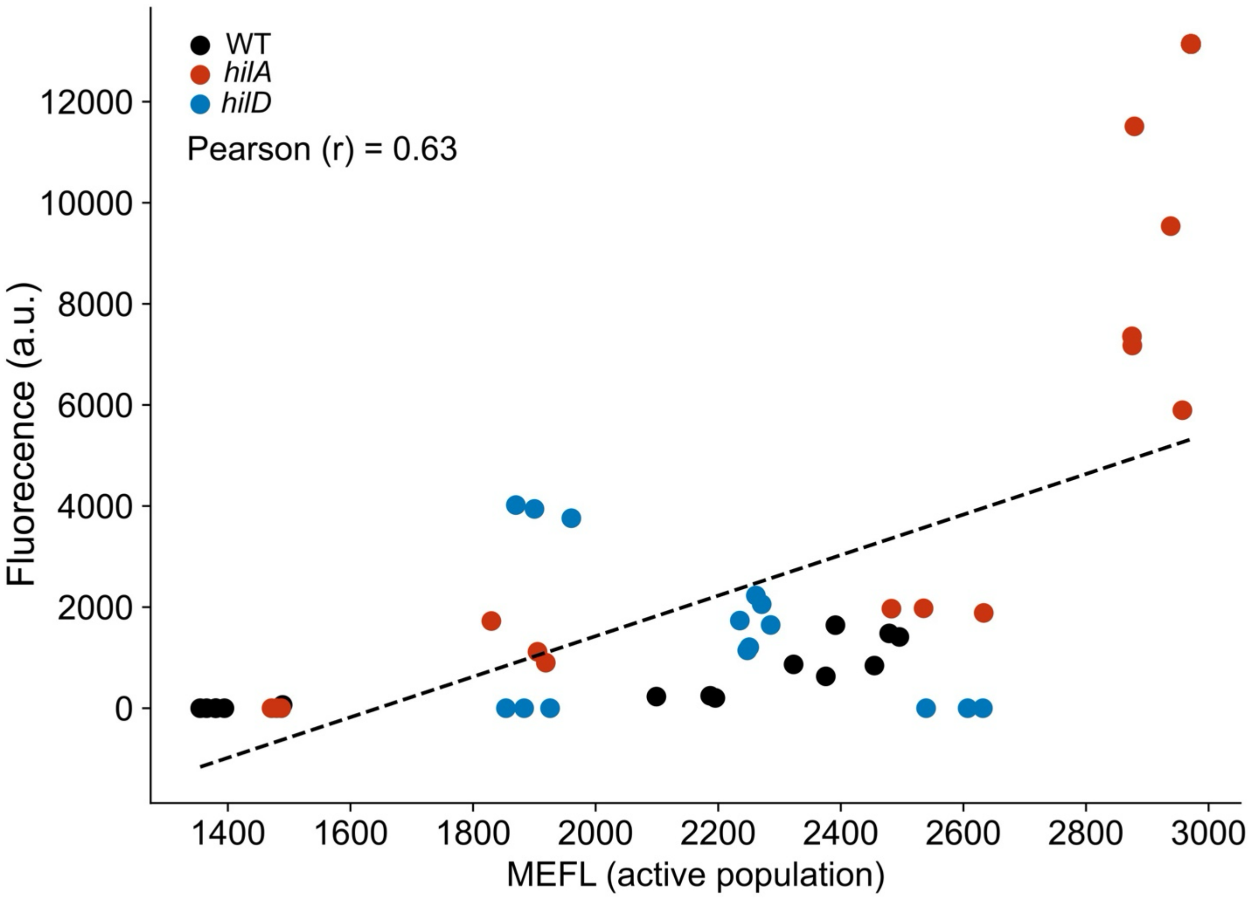


**Supplemental Figure 3:** Scatterplot of assembly measurement via flow cytometry (MEFL, active population) and mFAP secretion fluorescence (a.u.) over time for the three conditions tested: a WT *S*. Typhimurium strain, or *S*. Typhimurium strains overexpressing plasmid-encoded *hilA* or *hilD* to activate the T3SS. The black dotted line indicates the line of best fit.

| Metric | SR-SIM | Flow Cytometry |
|---|---|---|
| Samples per experiment | 15 | 60 |
| Sample preparation time | 4 days | 1 day |
| Data variation (coefficient of variation, CoV) | 44.3 | 3.3 |
| Data analysis time | 6 hr | 30 min |

**Supplemental Table 1:** Efficiency comparisons between flow cytometry and SR-SIM.
