## Supplemental Tables for "A high throughput flow cytometry assay for quantifying type 3 secretion system assembly in *Salmonella*"

**Table 1.1 Strains Used in This Work**

| Strain name | Organism | Description | Reference |
|---|---|---|---|
| ASTE13 | *Salmonella enterica* subsp. *enterica* serovar Typhimurium | High-secreting LT2-derived strain, similar to DW01 | [1] |
| ASTE13 Δ*prgI* | *Salmonella enterica* subsp. *enterica* serovar Typhimurium | Lacks T3SS needle monomer; secretion incompetent | [2] |
| ASTE13 *sipD::sipD-3xFLAG; prgI::cat-sacB* | *Salmonella enterica* subsp. *enterica* serovar Typhimurium | Genomically incorporated 3xFLAG sequence on the C-terminus of *sipD*; *cat-sacB* genomic insertion at the *prgI* locus | This study |
| ASTE13 *sipD::sipD-3xFLAG* | *Salmonella enterica* subsp. *enterica* serovar Typhimurium | Genomically incorporated 3xFLAG sequence on the C-terminus of *sipD* | This study |
| ASTE13 *prgI::6xHis-prgI* | *Salmonella enterica* subsp. *enterica* serovar Typhimurium | Genomically incorporated hexa histidine epitope on the N-terminus of *prgI* | [3] |
| ASTE13 *sipD::sipD-3xFLAG*; *prgI*::*6xHis-prgI* | *Salmonella enterica* subsp. *enterica* serovar Typhimurium | Genomically incorporated 3xFLAG sequence on the C-terminus of *sipD;* Genomically incorporated hexa-histidine epitope on the N-terminus of *prgI* | This study |
| ASTE13 *sipD*::*sipD-3xFLAG*; *prgI*::*6xHis-prgI-S49R* | *Salmonella enterica* subsp. *enterica* serovar Typhimurium | Genomically incorporated 3xFLAG sequence on the C-terminus of *sipD;* Genomically incorporated hexa- | This study |

| | | |
|---|---|---|
| | | histidine epitope on the N-terminus of *prgI* and S49R point mutation |
| DH10B | *Escherichia coli* | |

**Table 1.2 Plasmids Used In This Work**

| Plasmid name | ORFs Under Control | ORI | Antibiotic Resistance | Reference |
|---|---|---|---|---|
| PsicA-SicP-SptP-mFAP2a-2xFLAG-6xHis | *sicP; sptp-mFAP2a-2xFLAG-6xHis* | colE1 | Chloramphenicol | [4,5] |
| PlacUV5-hilA | *hilA* | p15a | Kanamycin | [2] |
| PlacUV5-hilD | *hilD* | p15a | Kanamycin | [6] |
| pSIM6 | *gam, beta, exo* | pSIC101 | Ampicillin, Carbenicillin | [7] |

**Table 1.3 Primers Used In This Work**

| Primer Sequence | Amplicon | Used For: |
|---|---|---|
| AGGCCATTGGTATTT CCCAAGCCCACTTTA ATTTAACGTAAATAA GGAAGTCATT ATCAAAGGGAAAAC TGTCCATAT | *Cat-sacB* | Forward primer for amplifying *catsacB* from the *E. coli* TUC01 genome with *prgI* locus homology arms |
| TAACGGCATTCTCAG GGACAATAGTTGCA ATCGACATAATCCAC CTTATAACTGA TGTGACGGAAGATC ACTTCG | *Cat-sacB* | Reverse primer for amplifying *catsacB* from the *E. coli* TUC01 genome with *prgI* locus homology arms |
| CCCAAGCCCACTTTA ATTTAACGTAAATAA GGAAGTCATTATGGC AACACCTTGGTCAGG | $prgI^{S49R}$ | Forward primer for amplifying $prgI^{S49R}$ and incorporating into *prgI* locus |

| | | |
|---|---|---|
| CCCTATAACGGCATT CTCAGGGACAATAGT TGCAATCGACATAAT CCACCTTATAACTGA | *prgI*$^{S49R}$ | Reverse primer for amplifying *prgI*$^{S49R}$ and incorporating into *prgI* locus |
| CCCAAGCCCACTTTA ATTTAACGTAAATAA GGAAGTCATTATGCA CCACCACCACCACCA CGCAACACCTTGGTC AGG | *prgI-6xHis* | Forward primer to add N-terminal 6xhis tag to *prgI* and *prgI* mutant |
| TGTGATTTAATCGCGCT CCTGATGGCGAACTGG GGATATTTGTGACGGA AGATCACTTCG | *Cat-sacB* | Forward primer for amplifying *catsacB* from the *E. coli* TUC01 genome with *sipD* locus homology arms |
| GAGTCCTTACACTTGTA ACCATTATTAATATCCT CTTCTGATCAAAGGGAA AACTGTCCATAT | *Cat-sacB* | Reverse primer for amplifying *catsacB* from the *E. coli* TUC01 genome with *prgI* locus homology arms |
| TGTGATTTAATCGCGCT CCTGATGGCGAACTGG GGATATTATGCTTAATA TTCAAAATTATTCCGCT TCTCC | *sipD-3xFLAG* | Forward primer for amplifying *sipD-3xFLAG* and incorporating into *sipD* locus |
| GAGTCCTTACACTTGTA ACCATTATTAATATCCT CTTCTGTTATCCTTGCA GGAAGCTTTTGG | *sipD-3xFLAG* | Reverse primer for amplifying *sipD-3xFLAG* and incorporating into *sipD* locus |
